# The framework of plant regeneration in duckweed (*Lemna turonifera*) comprises genetic transcript regulation and cyclohexane release

**DOI:** 10.1101/2021.07.22.453434

**Authors:** Lin Yang, Jinge Sun, Jie Yao, Yaya Wang, Congyu Yan, Junyi Wu, Qiuting Ren, Ling Zhao, Jinsheng Sun

## Abstract

Regeneration is important for vegetative propagation of excellent variety, detoxification and the obtain of transgenic plant, but plant regeneration is time-consuming. Here, we found that duckweed regeneration could be enhanced by regenerating callus. Firstly, Genetic transcript regulation has been applied to study the molecular mechanism controlling regeneration. Auxin related genes have been significantly down-regulated in regenerating callus. Cytokinin signal pathway genes have been up-regulated in regenerating callus. Secondly, volatile organic compounds release has been analysised by gas chromatography/mass spectrum during the stage of plant regeneration, and 11 kinds of unique volatile organic compounds in the regenerating callus were increased. Among them, cyclohexane treatment enhanced duckweed regeneration by initiating root. Moreover, Auxin signal pathway genes were down-regulated in callus treated by cyclohexane. All together, these results provide novel mechanistic insights into how regenerating callus promotes duckweed regeneration.

**Graphical abstract:** 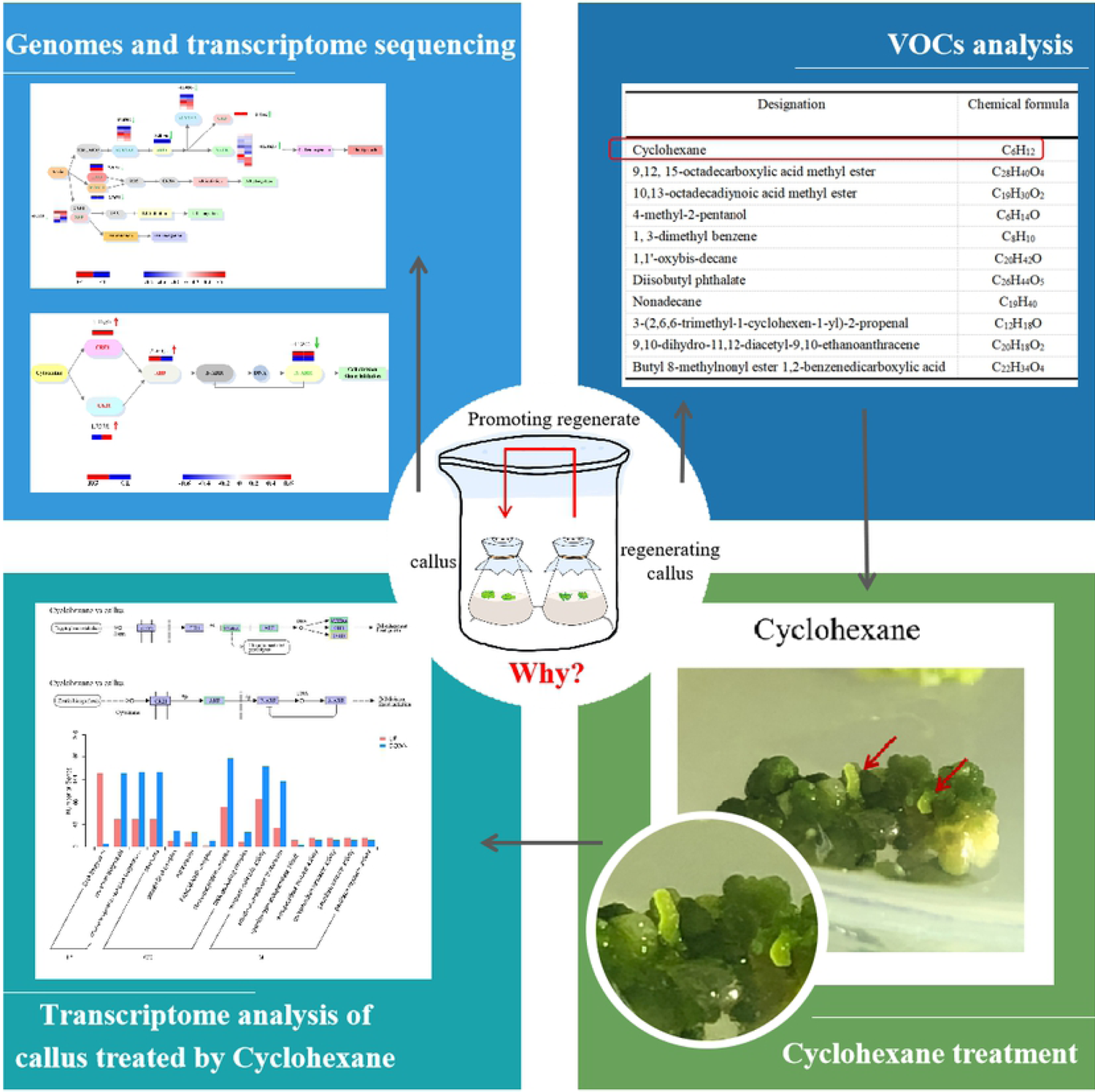

## Introduction

Regeneration of entire plants from callus in vitro depends on pluripotent cell mass, which provides rise to a new organ or even an entire plant^1–2^. Regeneration was widely used for vegetative propagation of excellent variety, detoxification and the obtain of transgenic crops^3–4^. A large number of studies have focused on the molecular framework of de novo organ formation in *Arabidopsis thaliana*. The molecular factors of cellular pluripotency during the regeneration of plants have been investigated thoroughly. However, the regulatory modules in monocot plants were little in-depth study. Duckweed, with the advantages of fast reproduction, high protein content^5^, and distinguished tolerance for a variety of toxic substances^6–7^, has been applied as a monocotylous modle plant for gene-expression systems. In duckweed, stable transformation mediate by *Agrobacterium* depends on efficient callus regeneration protocols.

Here, we use transcriptome sequencing technology to explore the molecular mechanism of plant hormones regulating callus regeneration^8^. Nevertheless, there is no study focus on the transcriptome analysis during the regeneration in duckweed. In former studies, it has been reported that the growth and development of callus was mediated by many plant hormones^5^. The balance of auxin and cytokinin is the basis for vitro tissue culture^9^. Explants can be incubate to callus on auxin-rich callus-inducing medium (CIM). And on cytokinin-rich shoot inducing medium (SIM), the vigorous callus can be induce to novo shoots. It is emergent to study the mechanism of duckweed regeneration via dynamic hormonal and transcriptional changes.

The volatile organic compounds (VOCs) could be produced to defense against herbivores, and it may also play a secondary role in attracting natural enemies, which is allelopathy^10–11^. For example, the VOCs of *Artemisia frigida Willd* play an allelopathic role on the seed germination of pasture grasses^12^. Does allelopathy play a role during plant regeneration? Interestingly, we found the plant regeneration could be promoted by regeneration callus. Why? The global insight on the signal and VOCs released from regenerating callus needs to be investigated.

Here, the main objectives has been studied: (i) the molecular mechanism controlling regeneration by comprehensive transcriptomic comparison between callus and regenerating callus; (ii) which VOCs have been increased during the stage of plant regeneration; (iii) the allelopathic effects of VOCs on the inducement of callus regeneration; (iv) the transcriptome analysis on the regenerating callus which has been promoted by VOCs.

## Results

### Promoted effect of regenerating tissue

Frond regeneration of duckweed has been promoted when co-cultured with regenerating callus (Co). Frond formed in 14 d with Co treatment, and duckweed regenerated at 21 days with with Co treatment (Fig 1a). In Co group, significant enhancement was found in the percentage of callus regeneration (77.3 %). Compared with that, the callus regeneration percentage without co-culture was 53.6% (Fig 1b). Thus, the callus regeneration has been significantly increased by Co treatment. Fig. 1 The co-cultured of callus and regenerating callus.

**Fig. I.**
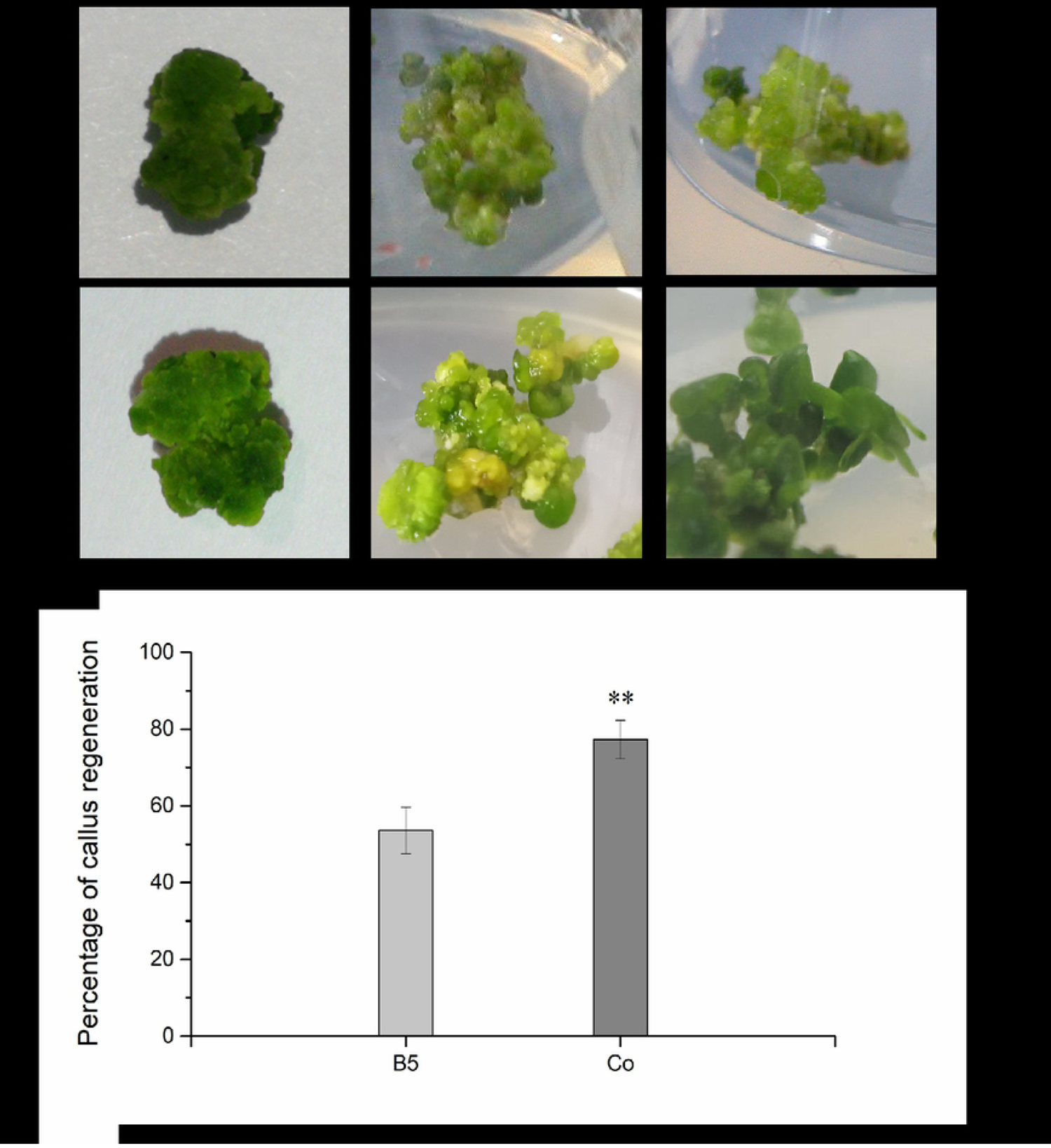
The co-cultured of callus and regenerating callus. a The large beaker was sealed with plastic wrap and perforated with a sterile toothpick. b The ratio of callus regeneration between the control group (85) and co-culture condition.

### Transcriptome analysis identifies Genes and Genomes (KEGG) and differentially expressed genes (DEGs) in regenerating callus

To compare the enriched pathways between regenerating callus (RG) and callus (CL), KEGG pathway analysis has been conducted (Fig. 2). The top 20 KEGG pathways with the highest representation of DEGs have been analyzed. We selected the 20 pathway items that were most significant in the enrichment process to be shown in this diagram. As shown in Fig. 2a, the “Photosynthesis antenna proteins” was the most significantly enhanced pathway in the top 20 up-regulated KEGG pathways with the highest Rich Factors of RG vs CL. This indicated that the expression of antenna protein increased after the callus developed into regenerated tissue. Antenna proteins were very important for plant photochemical reactions and could mediate the core of plant photosynthesis. The most significantly down-regulated pathway was the “Ribosome”, “Pyrimidine metabolism”, “Mismatch repaire”, “Homologous recombination”, “DNA replication” and “Base excision repair”, which were among the top list of enriched pathways (Fig. 2b), these were all related to the replication of DNA.

**Fig. 2.**
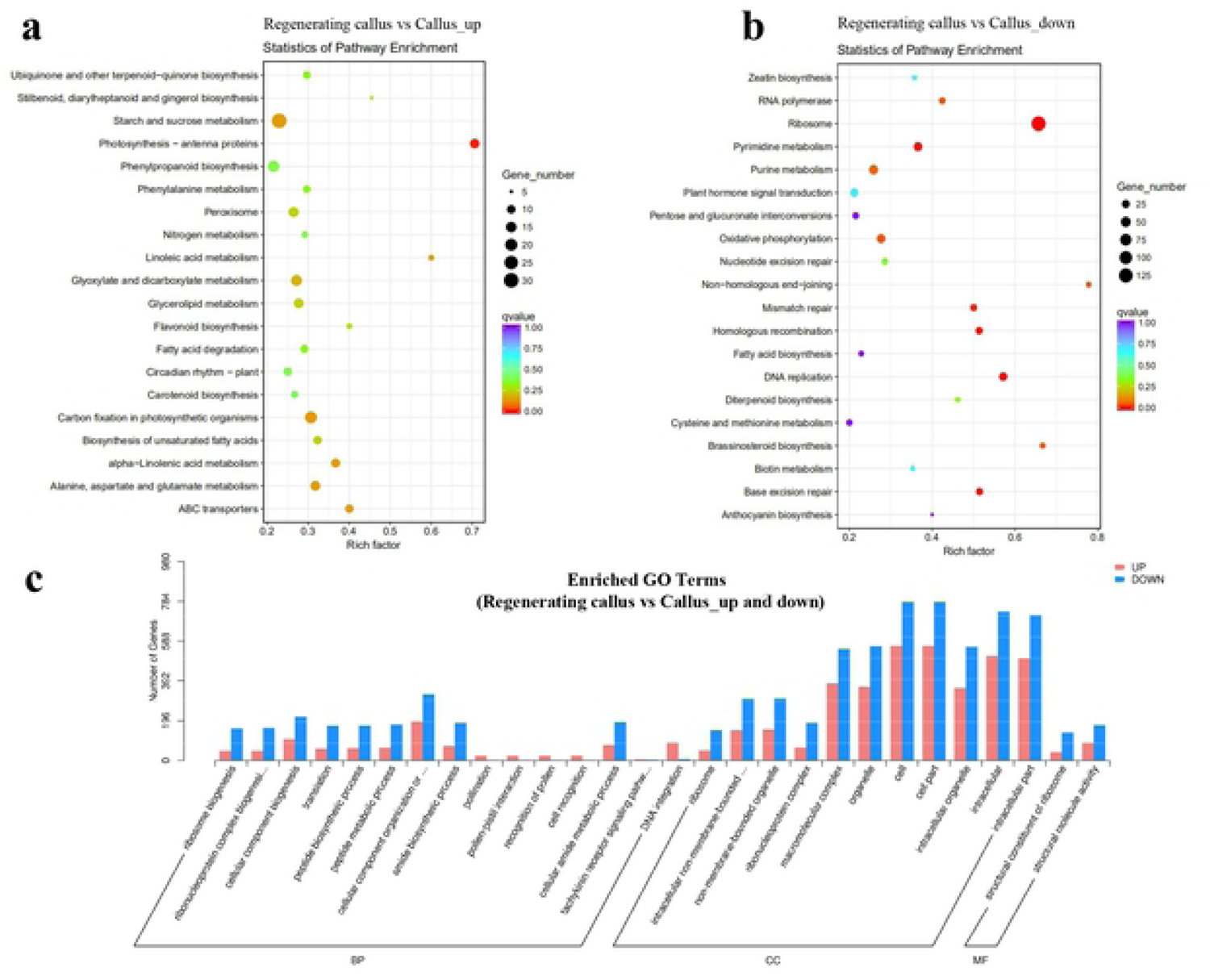
Statistic of KEGG pathway enrichrnent and the nurn ber of enriched genes in different gene ontolog y (GO) categories in RG vs CL. **a** The top 20 up KEGG pathways with the highest Rich Factors of RG vs CL, The KEGG Pathway enric h1nent hub diagra1n: The vertical axis represents pathway narne, the horizontal axis represents the Rich factor corresponding to pathway, and the colo ur of the dots represents the sizeof the Q value; the smaller the Q value, the closer the colour to red; the nurnber of different genes contained in each pathway is represented by the size of the dots, and the value range of qvalue was [0, 1], and the closer to zero, the more significant the enrichrnent; b The top 20 down KEGG pathways with the highest Rich Factors of RG vs CL; c GO terms associated with DEGs in RG and CL. The x-coordinate was GO the next level of the three categories GO entry, and ordinate was the number of different genes commented to the entrance.

In order to understand the difference of DEGs in the regenerating callus, gene ontology enrichment analysis was conducted in RG vs CL. As shown in Fig. 2c, “cell” “cellpart” and “intracellular” were in biological process with the most up-regulated and down-regulated DEGs. These were followed by “macromolecular complex” and “organelle” in the category of biological process with the most up-regulated and down regulated DEGs. “DNA integration”, “pollination”, and “cell recognition” were up-regulated DEGs, without down-regulated (Fig.2). Fig. 2 Statistic of KEGG pathway enrichment and the number of enriched genes in different gene ontology (GO) categories in RG vs CL.

### Expression changes of genes related to Auxin and root development in regenerating callus

The mRNA expression was conducted by Novogene in order to study the gene that participated during callus regeneration. The course of auxin signal pathway and related response factors have been described as Fig. 3. Transport inhibitor response 1 (TIR1) and stem cell factor (SCF), initiating subsequent signal transduction by binding of auxin, have been down-regulated in the regenerating callus. As a transcriptional activator, auxin response factor (ARF) could regulate auxin reaction by binding with auxin-responsive protein IAA (AUX/IAA). In this study, AUX/IAA and ARF have been down-regulated significantly, by 13.0309 and 3.0056 log^2^ Fold Change, respectivly. Auxin early response factor could be divided into three categories, which were AUX/IAA, Gretchen Hagen 3 (GH3) and small auxin-up RNA (SAUR). GH3 and SAUR have been down-regulated during regeneration, as well. ETHYLENE-RESPONSIVE FACTOR3 (ERF3) and WUSCHEL-RELATED HOMEOBOX 11 (WOX11), playing a role in the initiation and regulation of adventitious roots (ARs), were both down-regulated. Also, lateral roots (LRs) and root hairs (RHs) were rely on zinc finger protein (ZFP) and cytochrome P450 (CYP2). The expression of ZFP was decreased by 4.0368 log^2^ Fold Change.

**Fig. 3.**
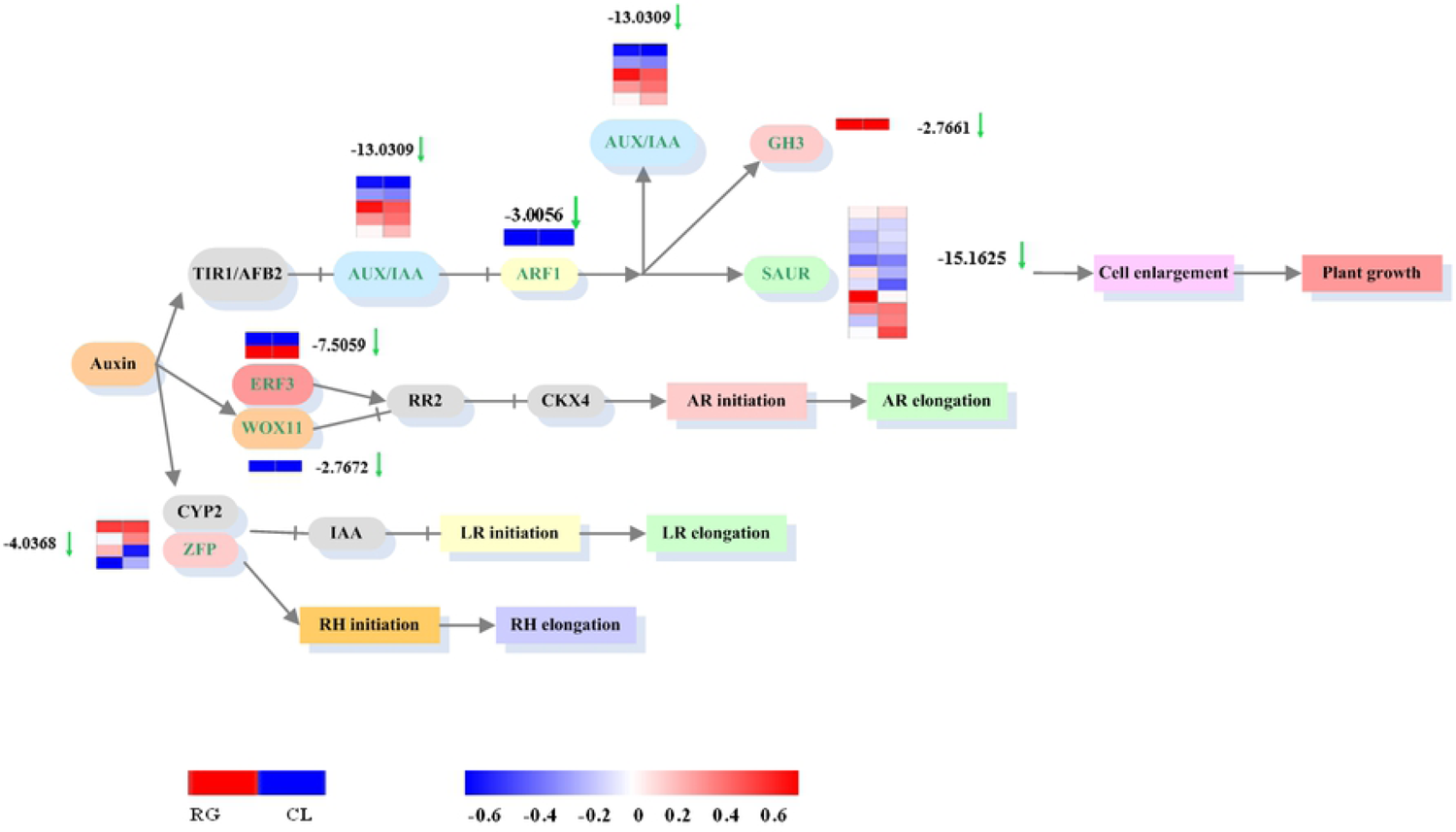
The comparison between regenerating callus and callus was related to auxin metabolism response and auxin signal transduction pathway. Arrows indicated the direction of processes, while red was up, green was down. As shown in the figure was auxin signal transduction, and various response factors were down-regulated. The color in this figure legend from red to blue, which meant log! 0 (FPKM+I) from high to low. Red meant high expression, blue meant low expression.

### Expression changes of genes related to cytokinins signal pathway in regenerating callus

To obtain candidates regulating regeneration, we studied the regulation of cytokinins signal pathway. Shown as Fig.4, cytokinin receptor 1(CRE1) and cytokinin independent 1(CKI1), as cytokinin receptors^13–14^, have been up-regulated in regenerating callus. Histidine phosphate transfer protein (AHP), interacting with CRE1 and CKI1, has been up-regulated by 2.9662 log ^2^ Fold Change. Type-A ARABIDOPSIS RESPONSE REGULATORS (A-ARR) plays a role as a negative feedback regulator, which inhibit the activity of type-B ARABIDOPSIS RESPONSE REGULATORS (B-ARR) and form a negative feedback cycle^15–16^. A-ARR has been down-regulated by 4.5266 log ^2^ Fold Change. It might be lead to overall up-regulated in cytokinins during the callus regenerating.

**Fig. 4.**
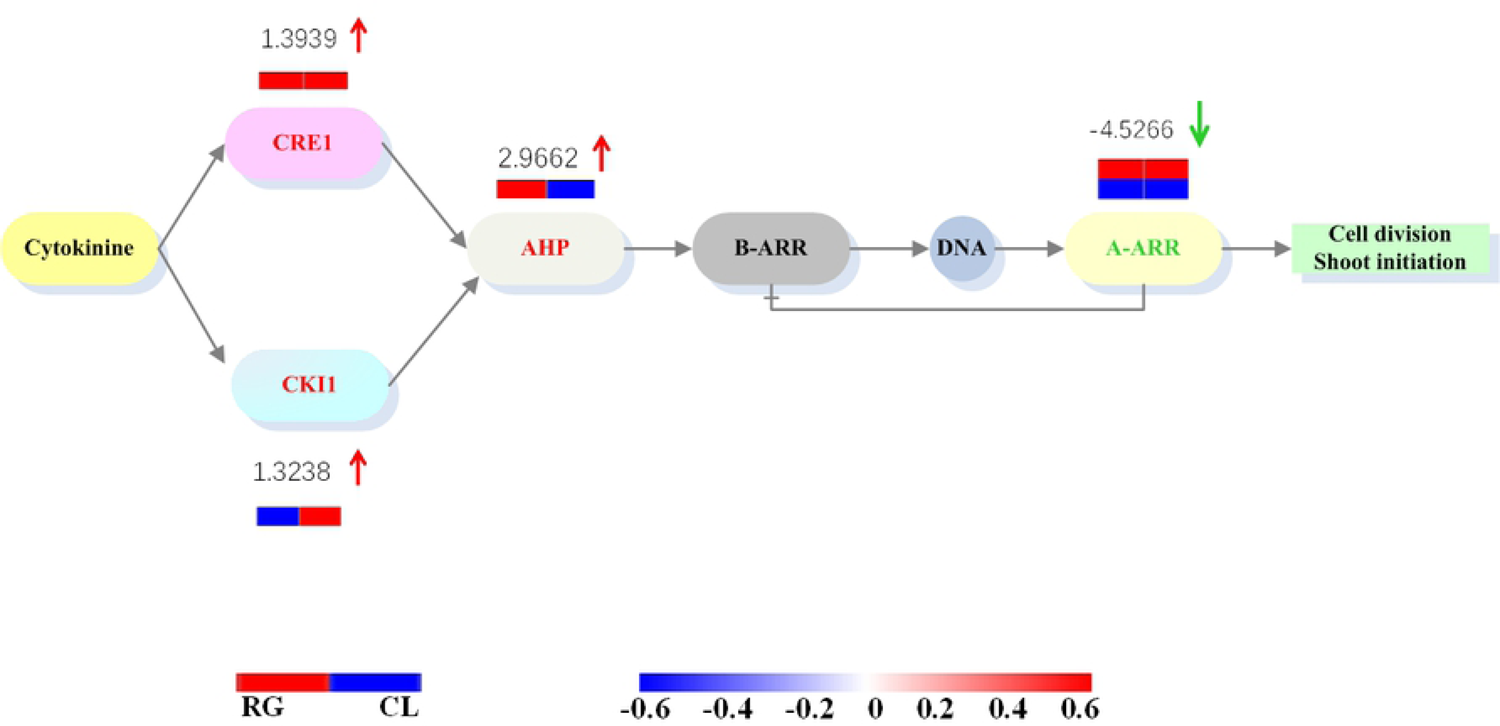
Comparing regenerating callus and callus was related to cytokinins metabolism response and cytokinins signal transduction pathway. Arrows indicated the direction of processes, while red was up, green was down. As shown in the figure was cytokinins signal transduction. CKII, CREI and AHP were up-regulated, but negative feedback regulator A-ARR was down-reg+ulated. The colour in this figure legend from red to blue, which meant loglO (FPKM+ l) from high to low. Red meant high expression, blue meant low expression.

### Changes of VOCs during callus regeneration

The VOCs of regenerating callus have been investigated. And the qualitative and quantitative analyses of the GC/MS data were obtained from NIST/EPA/NIH Mass Spectral Library, showed as Fig.5. Compared to the callus, 11 kinds of unique VOCs in the regenerating callus were enhanced (Table 1). The peak area of 1, 3-dimethyl benzene in the regenerating callus was 0.84*10^7^, 3.23 times than that in the callus. And the emission of 1, 3-dimethyl benzene increased the most in the regenerating callus. Besides, the content of 4-methyl-2-pentanol and cyclohexane also have been improved. Compared with the cyclohexane peak area of the callus (0.85*10^7^), the cyclohexane peak area of the regenerating callus was 1.28*10^7^, 4.3*10^6^ higher than that of callus. And the peak area of 4-methyl-2-pentanol was 2.1*10^7^, 2.33 times than that of callus.

**Fig. 5.**
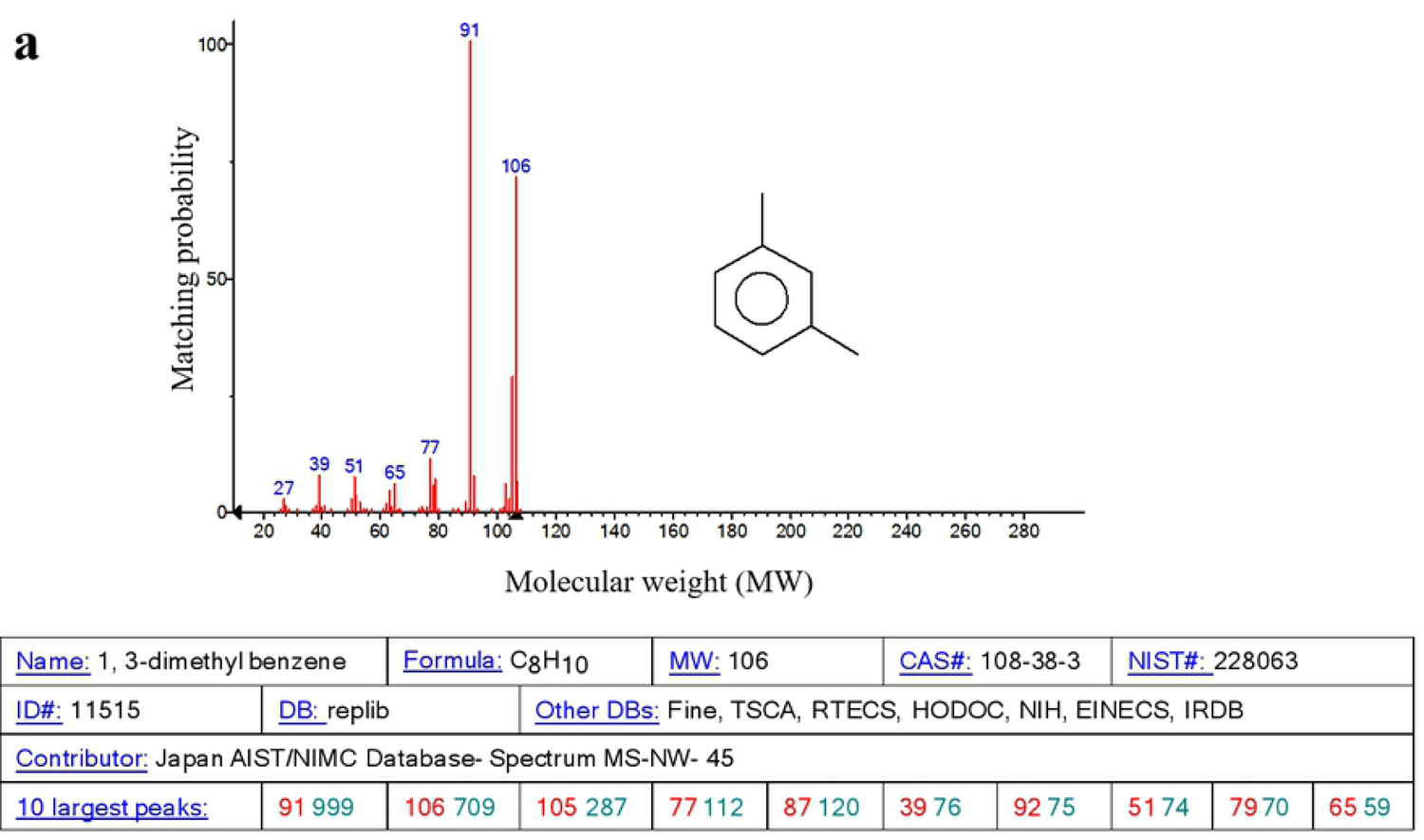
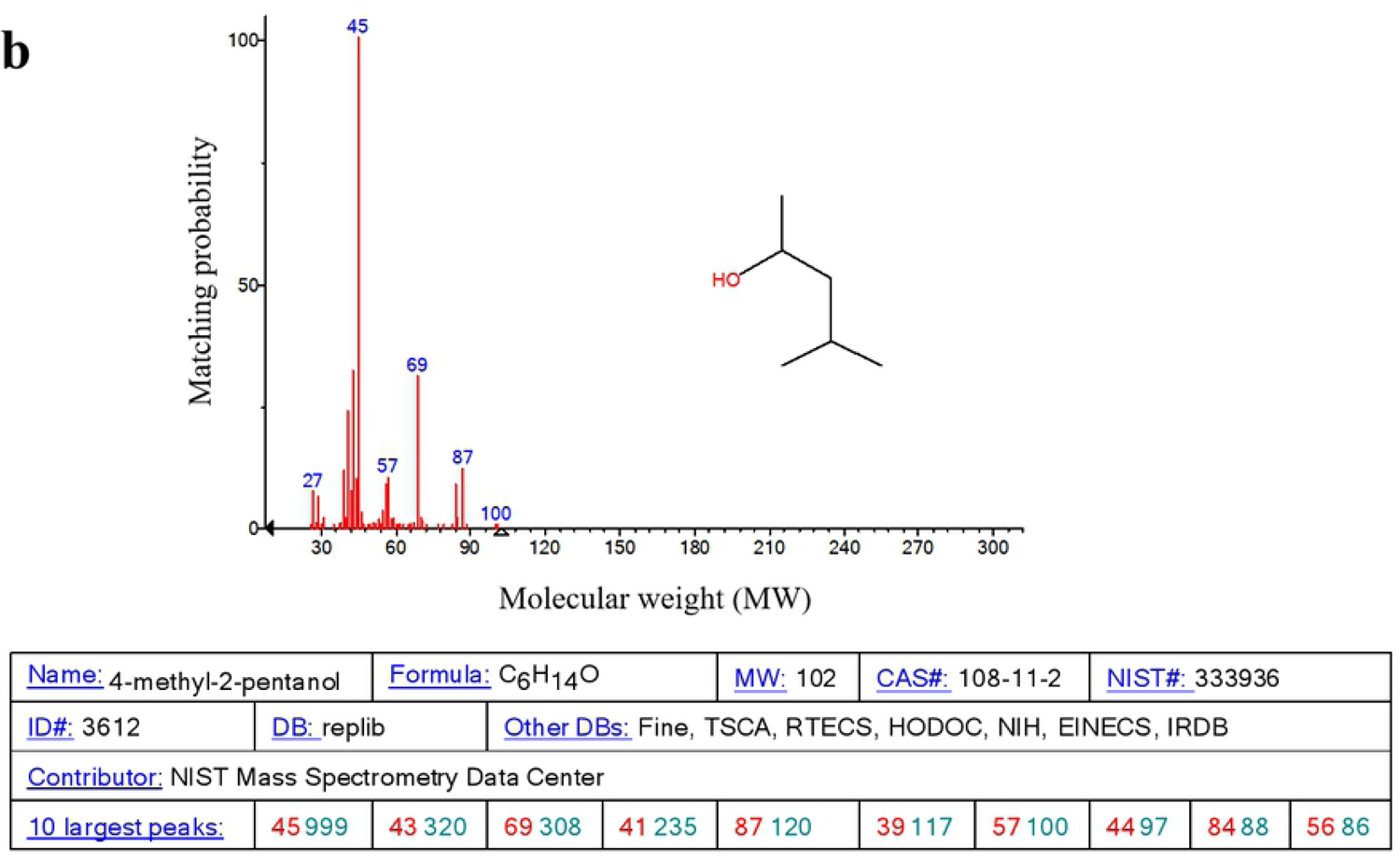
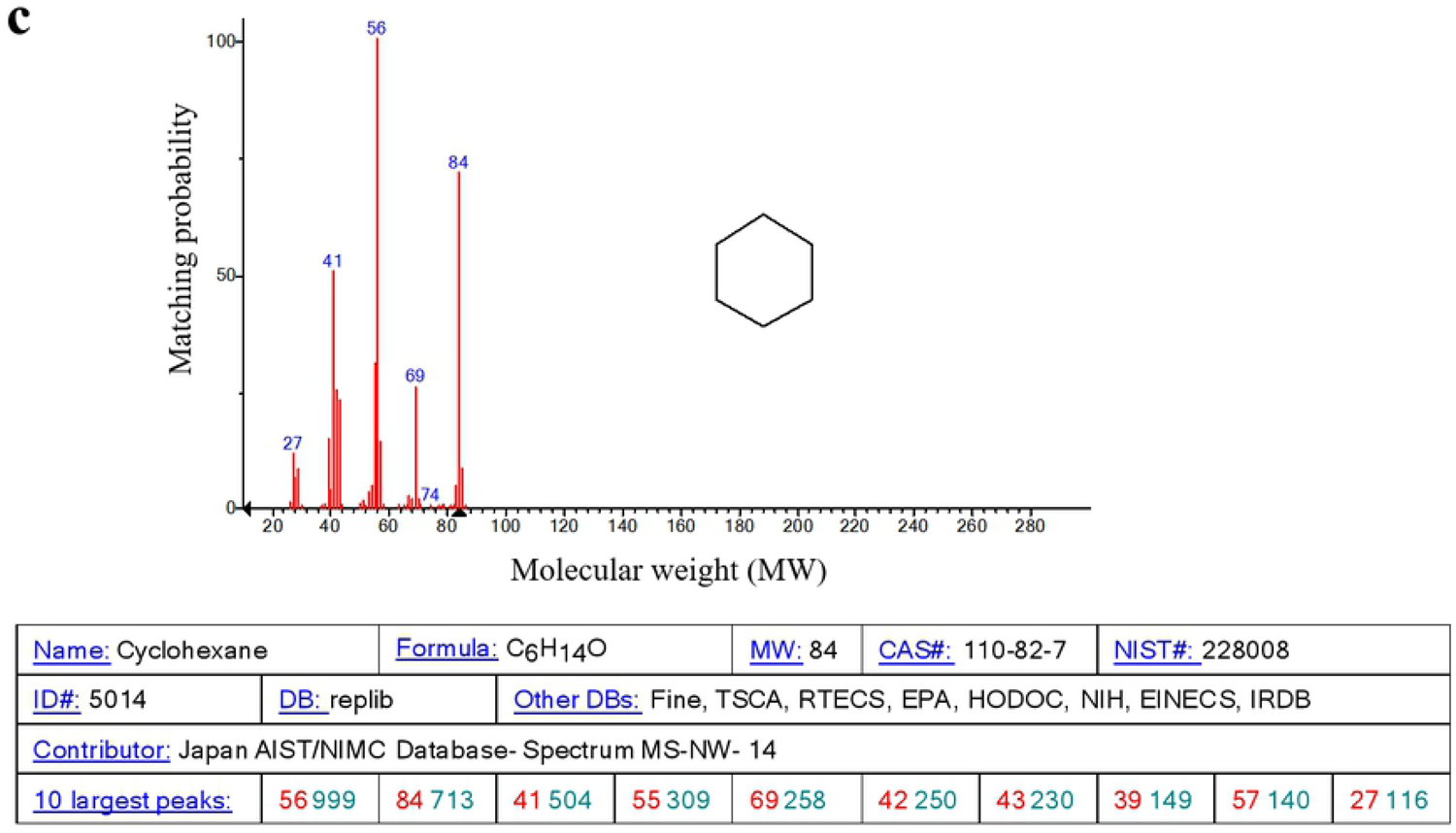
Three kinds of voes significantly up-regulated in the callus regeneration stage. The numbers in blue represented the mass-to-charge ratio (m/z) of a substance in the histogram. a Mass spectra of 1, 3-diinethyl benzene. b Mass spectra of 4-methyl-2-pentanol. c Mass spectra of cyclohexane.

**Table 1.**
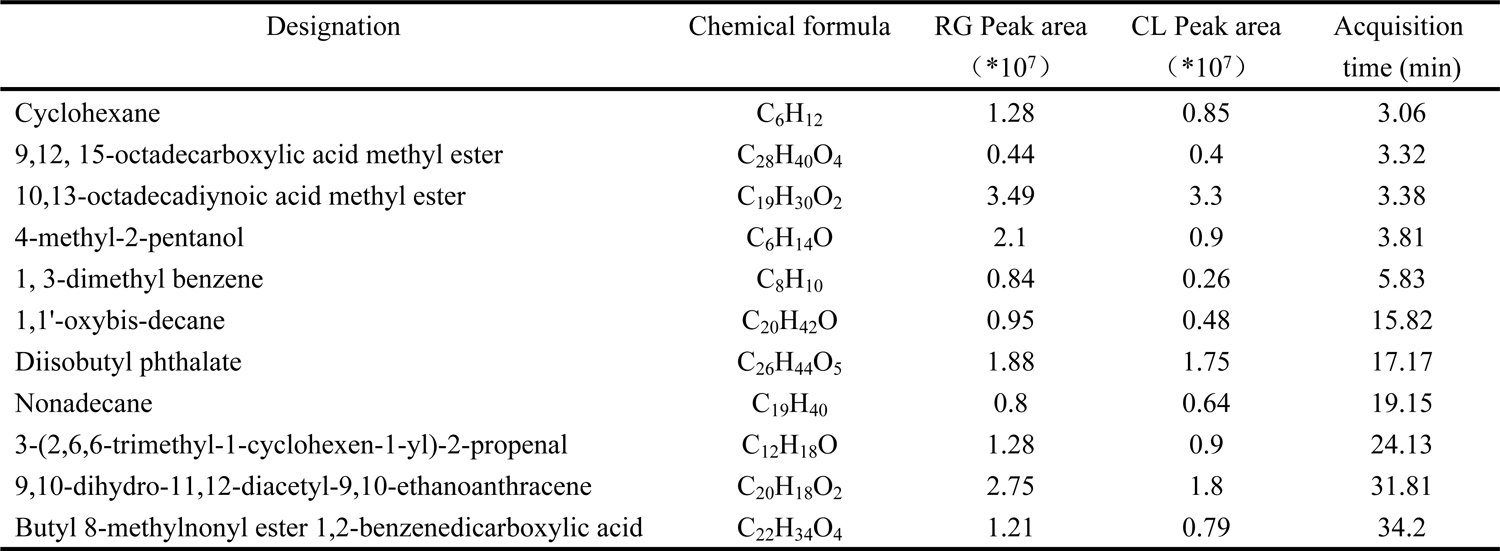
The main components of VOCs from regenerating callus and callus

### Callus regeneration was promoted by cyclohexane

In order to explore the effect of VOCs in callus regeneration, 1, 3-dimethyl benzene, 4-methyl-2-pentanol and cyclohexane were added to the medium of callus. As Fig. 6 showed, cyclohexane promoted the regeneration of callus significantly. After 16 days cyclohexane treatment, roots formed from the callus. The newborn roots could be distinctly observed shown as red arrow. However, 1, 3-dimethyl benzene and 4-methyl-2-pentanol groups have no obvious phenomenon of regeneration in 16 days. Fig. 6 Effects of 16 days’ treatment of callus by three VOCs (cyclohexane, 4-methyl-2-pentanol and 1, 3-dimethyl benzene).

**Fig. 6.**
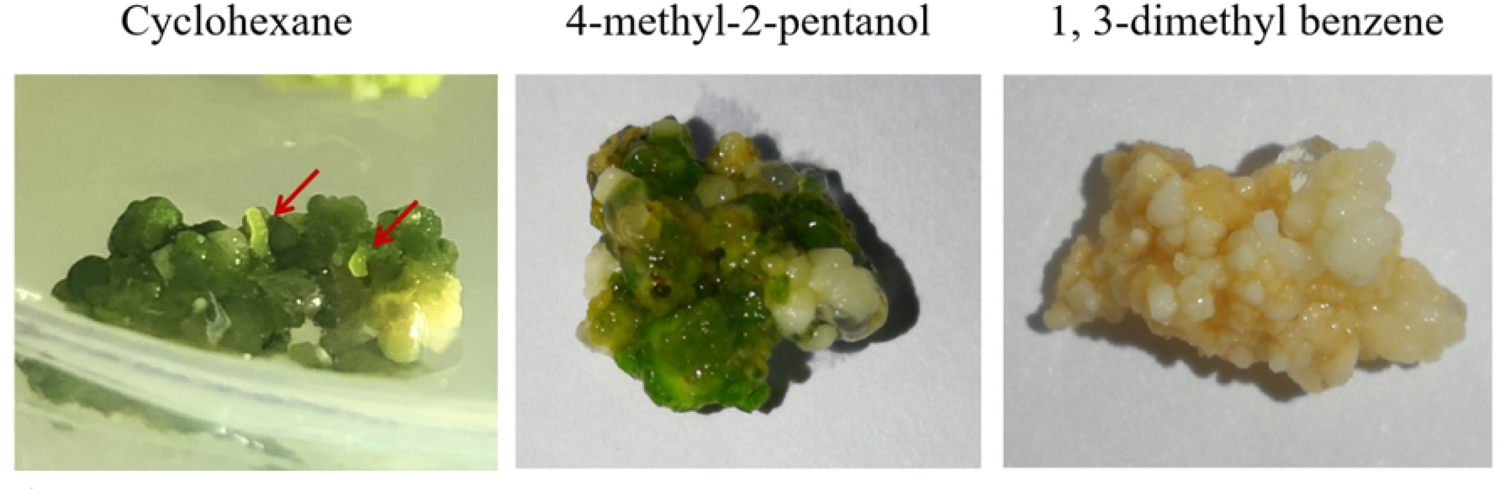
Effects of 16 days’ treatment of callus by three VOCs (cyclohexane, 4-methyl-2-pentanol and, I 3-dimethyl benzene).

### Transcriptome analysis identifies KEGGs and DEGs in callus treated by cyclohexane

Transcriptome analysis has been analyed to investigate the potential functions of KEGGs and DEGs in the callus treat hydrolyzing O-glycosyl compoundsed by cyclohexane. As shown in Fig. 7a, “RNA transport” and “glycolysis/gluconeoge, and galaclose metabolism” were in the biological process with the most down-regulated KEGGs. “Ribosome” was the the top-enriched pathway (Richfactor>0.55). It was followed by “photosynthesis”, and “oxidative phosphorylation” (Fig. 7b).

**Fig. 7.**
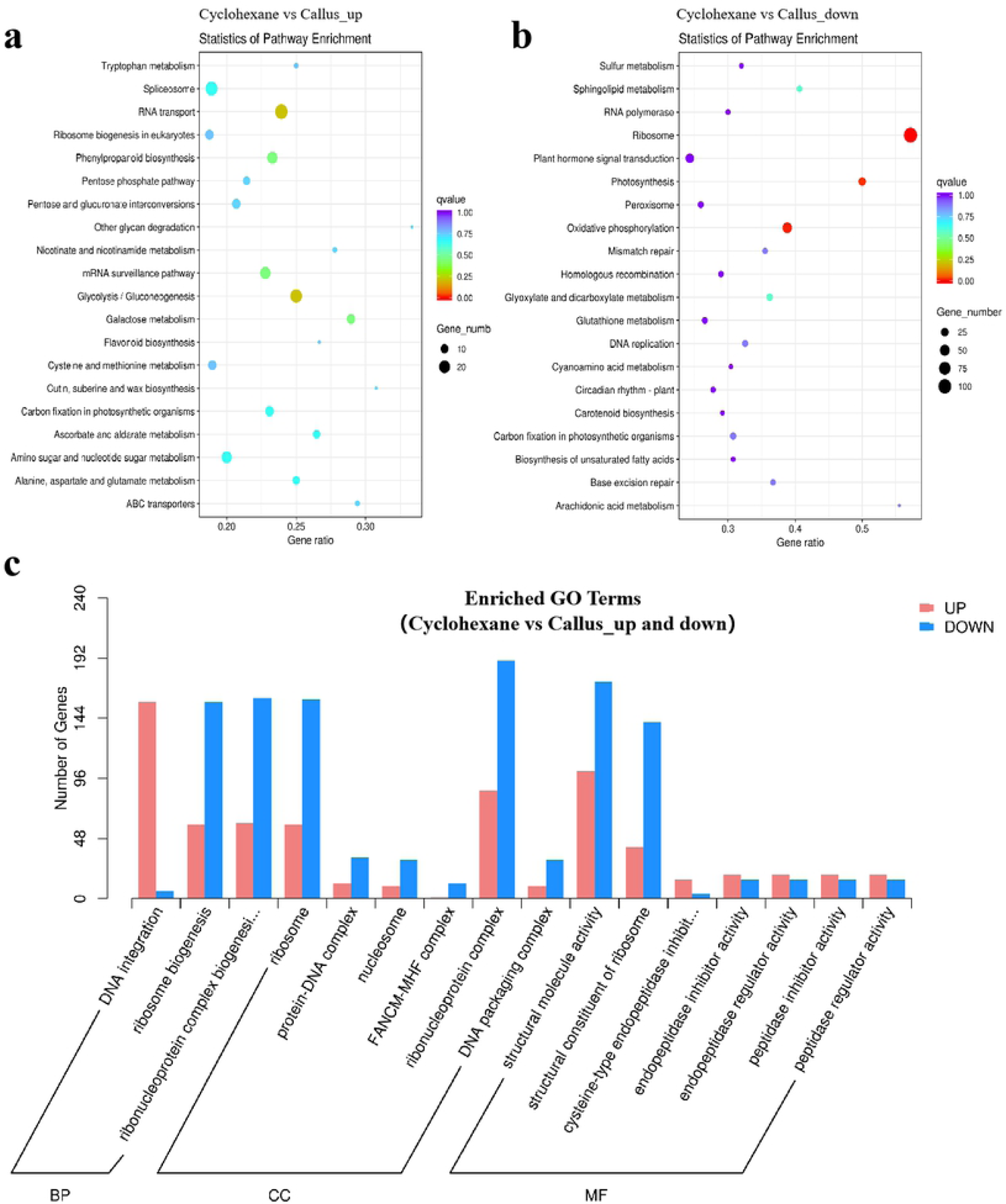
In the context of “Cyclohexane vs CL”, the top 20 KEGG pathways of up-regulated DEGs (a) and down-reg ulated DEGs (b) with the highest Rich Factors. GO terms associated with DEGs in “Cyclohexane vs CL”, the number of Enriched were up and down-regulated DEGs (c) in different gene ontology categories.

In order to understand the difference of DEGs in callus treated with cyclohexane, gene ontology enrichment analysis was conducted in callus treated by cyclohexane vs callus. As shown in Fig. 7c, “DNA integration”, “ribonucleoprotein complex” and “structrural molecule activity” were in biological process with the most up-regulated DEGs. These were followed by “ribosome biogenesis”, “ribonucleoprotein complex” and “ribosome” in the category of biological process with the most up-regulated DEGs. “ribonucleoprotein complex” and “structrural molecule activity” were were in biological process with the most down-regulated DEGs. (Fig. 7c).

### Comparsion of the expression of genes related to hormone in callus treated with cyclohexane and in the regenerating callus

In order to know molecular factors underlying the participation of hormone in callus regeneration, we first checked gene expression related to auxin signal pathway (Table 2). AUX/IAA and GH3 has been down regulated in both callus treated with cyclohexane and in the regenerating callus. A majority of SAUR have been down-regulated during regeneration and treated with cyclohexane (Fig. 8a). ERF3, cysteine-rich receptor and Zinc finger has been down-regulated as well.

**Fig. 8.**
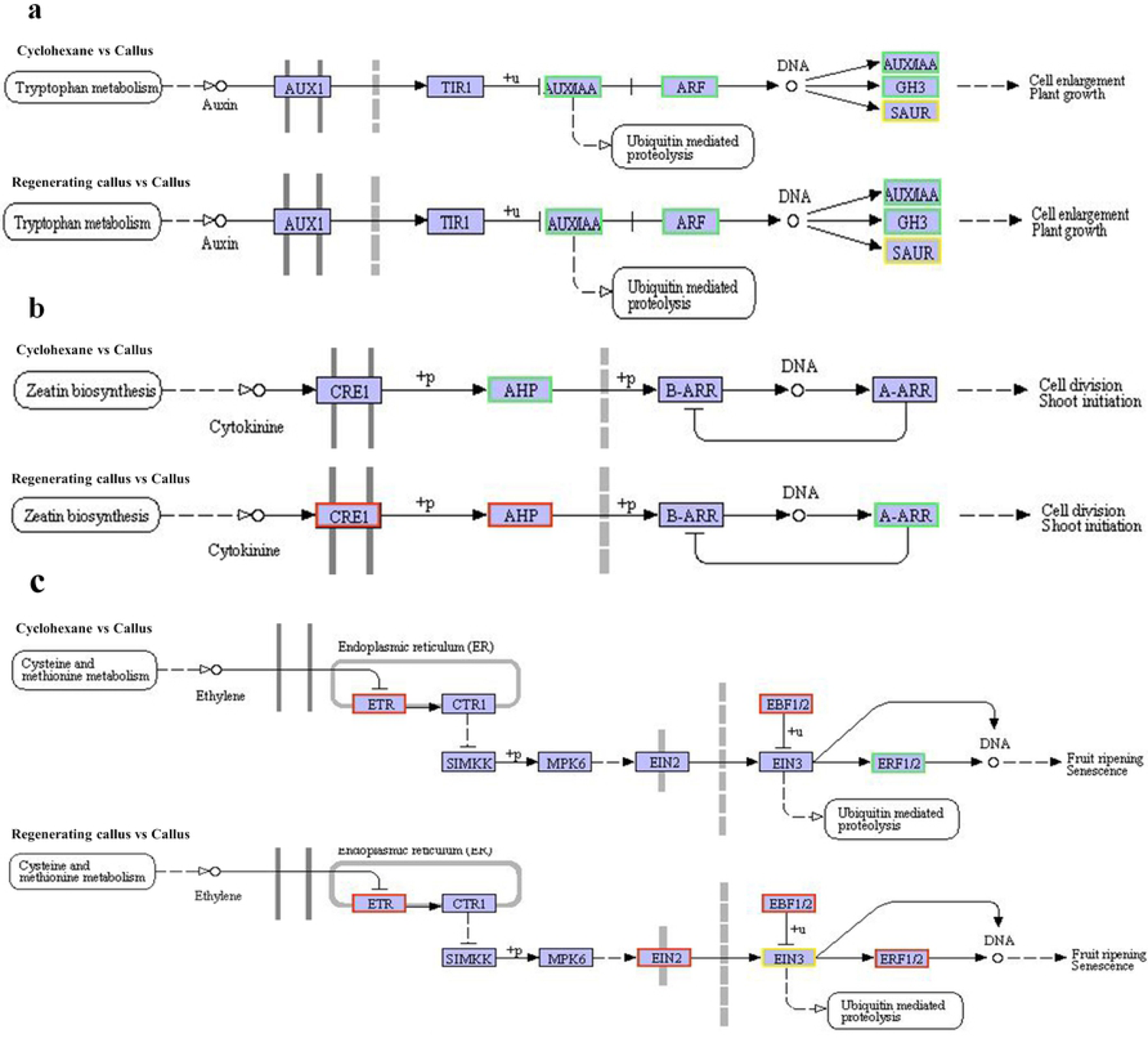
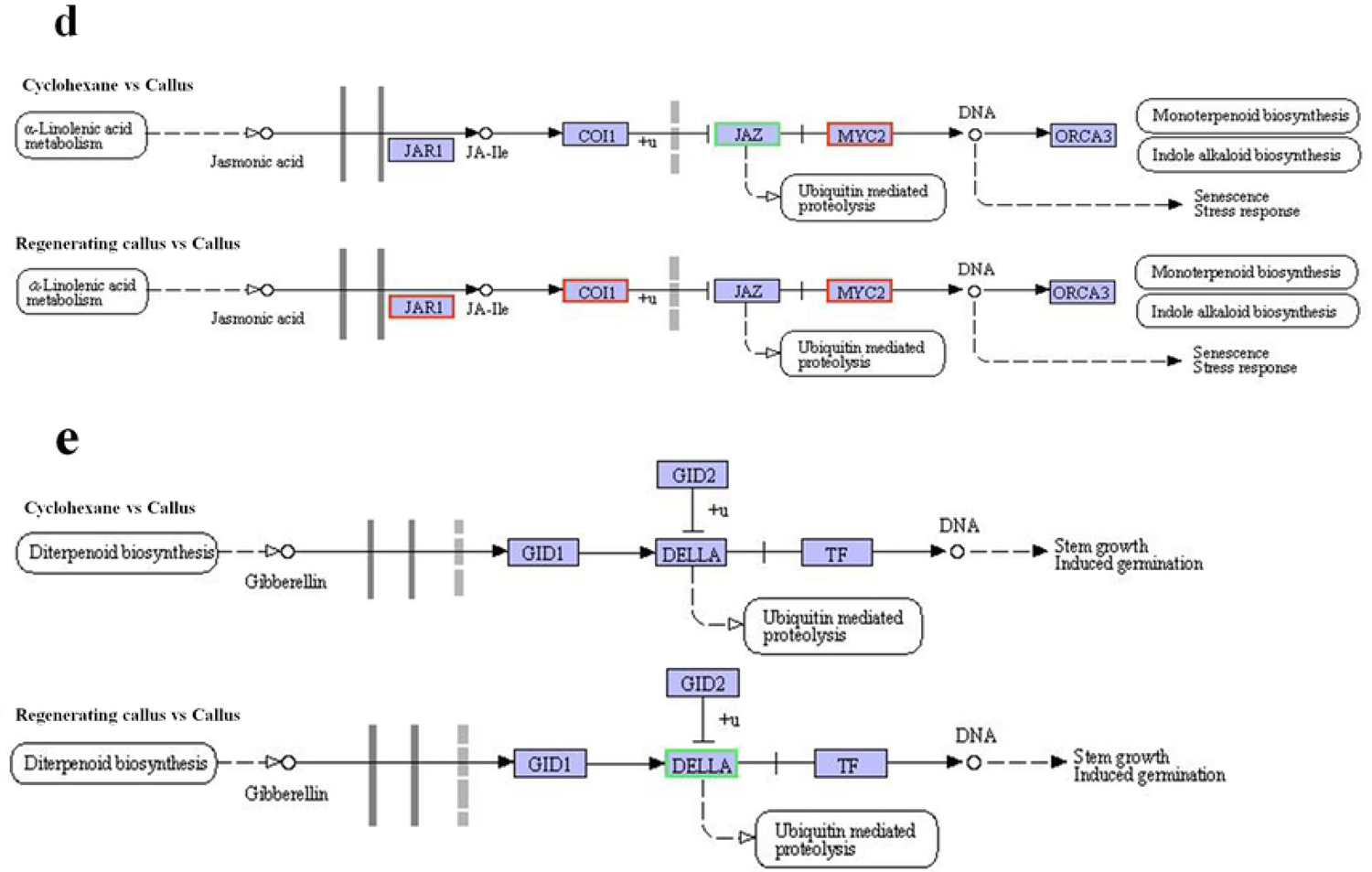
The pathway of biosynthesis of five types of plant hormone. Red meant high expression, and blue meant low expression. a The changes of genes in Auxin between regenerating callus and cyclohexane treatment callus. b The changes of genes in cytokinin between regenerating callus and cyclohexane treat1ne nt callus. c The differences of genes in ethylene between regenerating callus and cyclohexane treatn1ent callus. d The changes of genes injasmonic acid between regenerating callus and cyclohexane treatment callus. e The changes of genes in gibberellin between regenerating callus and cyclohexa ne treat1nent callus.

**Table 2.**
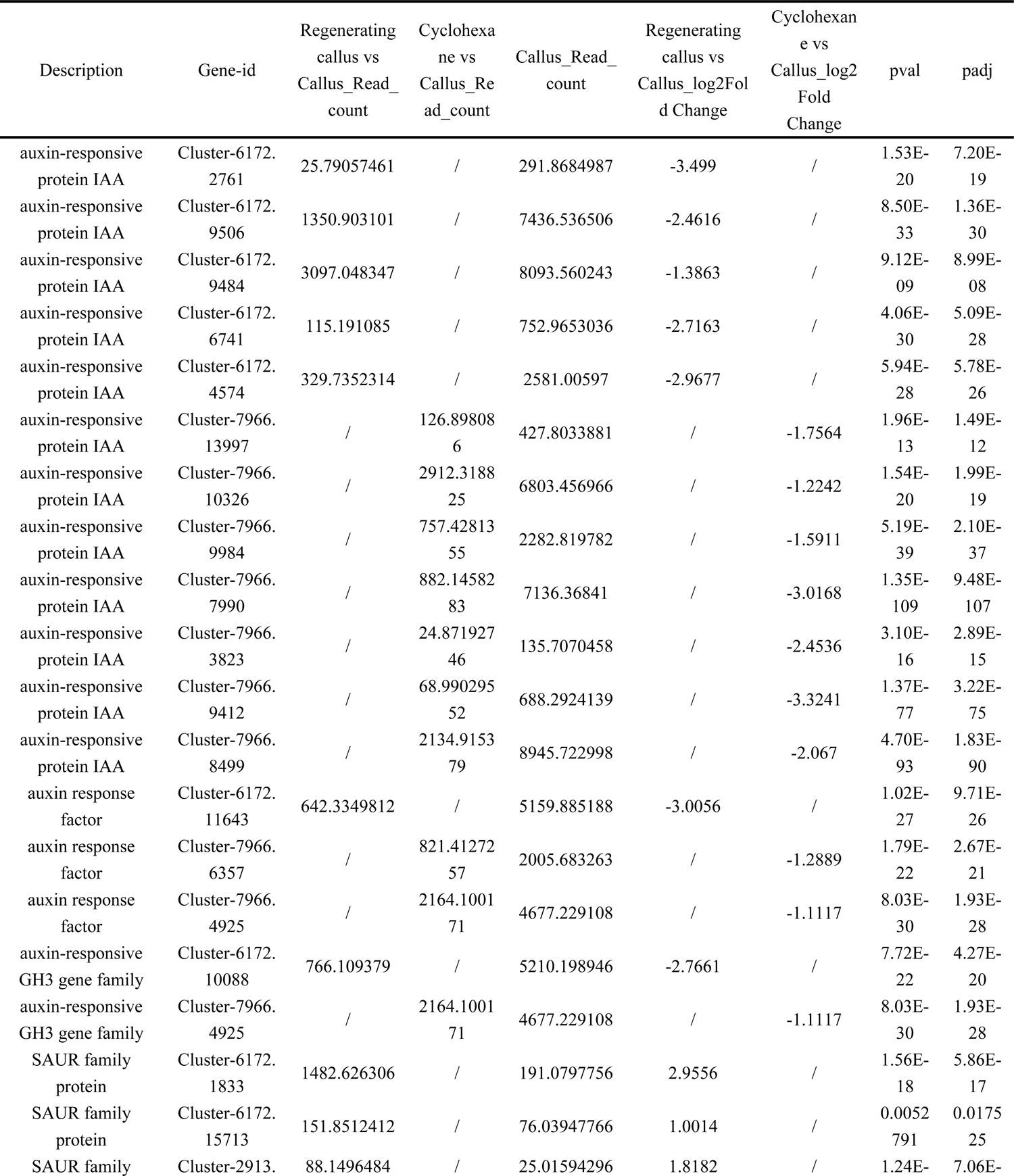
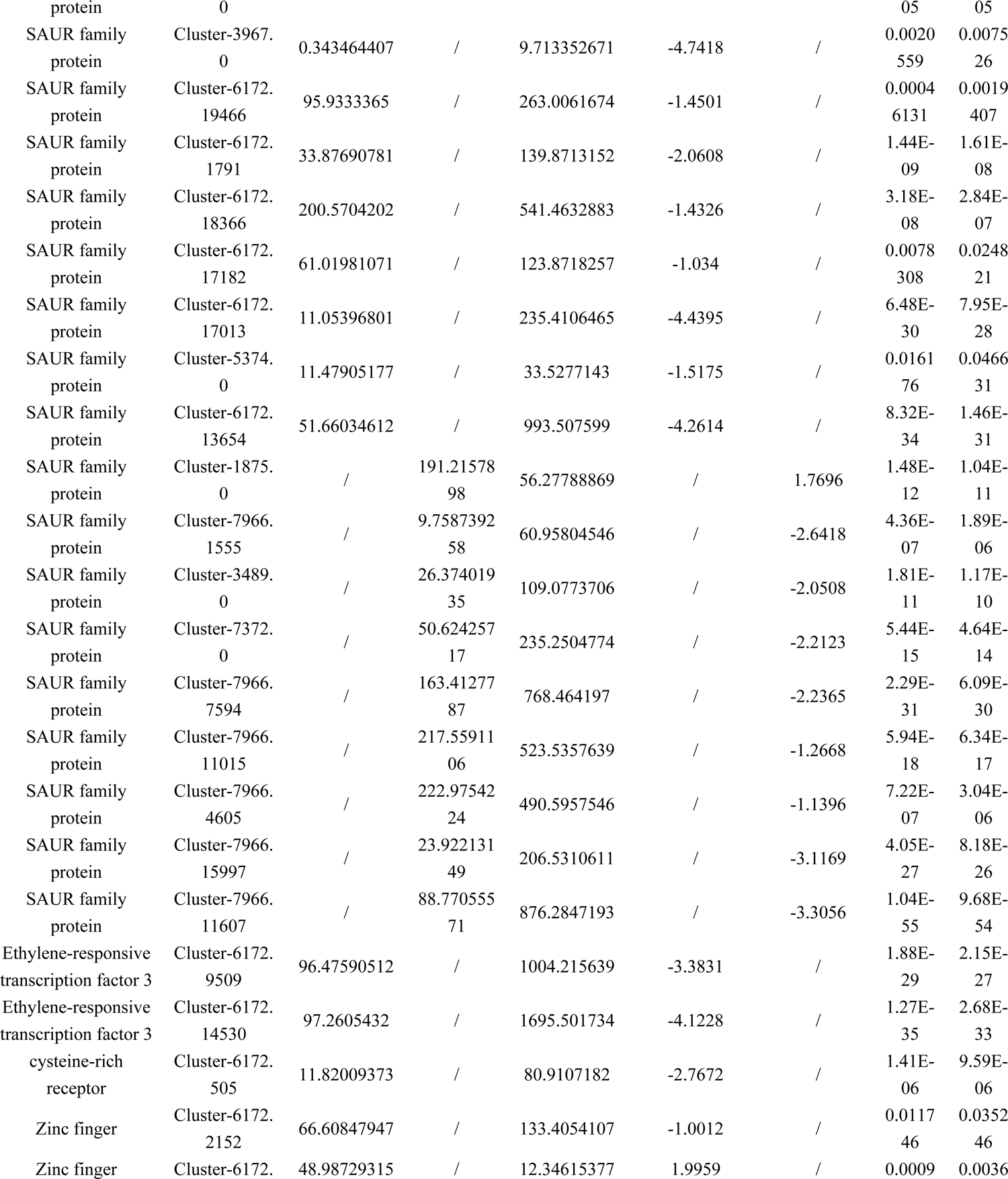
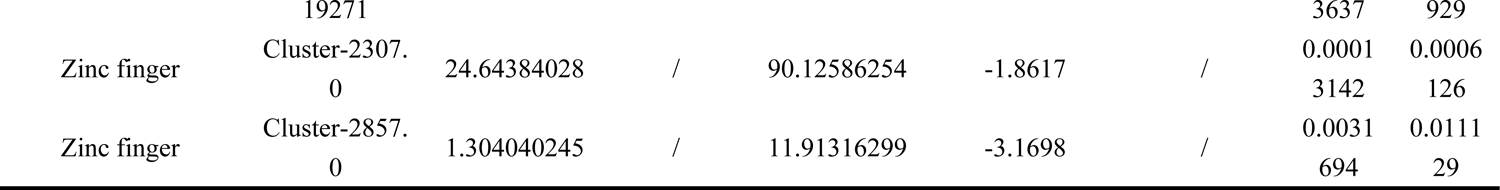
Gene expression in plant regeneration of Auxin

**Table 3.**
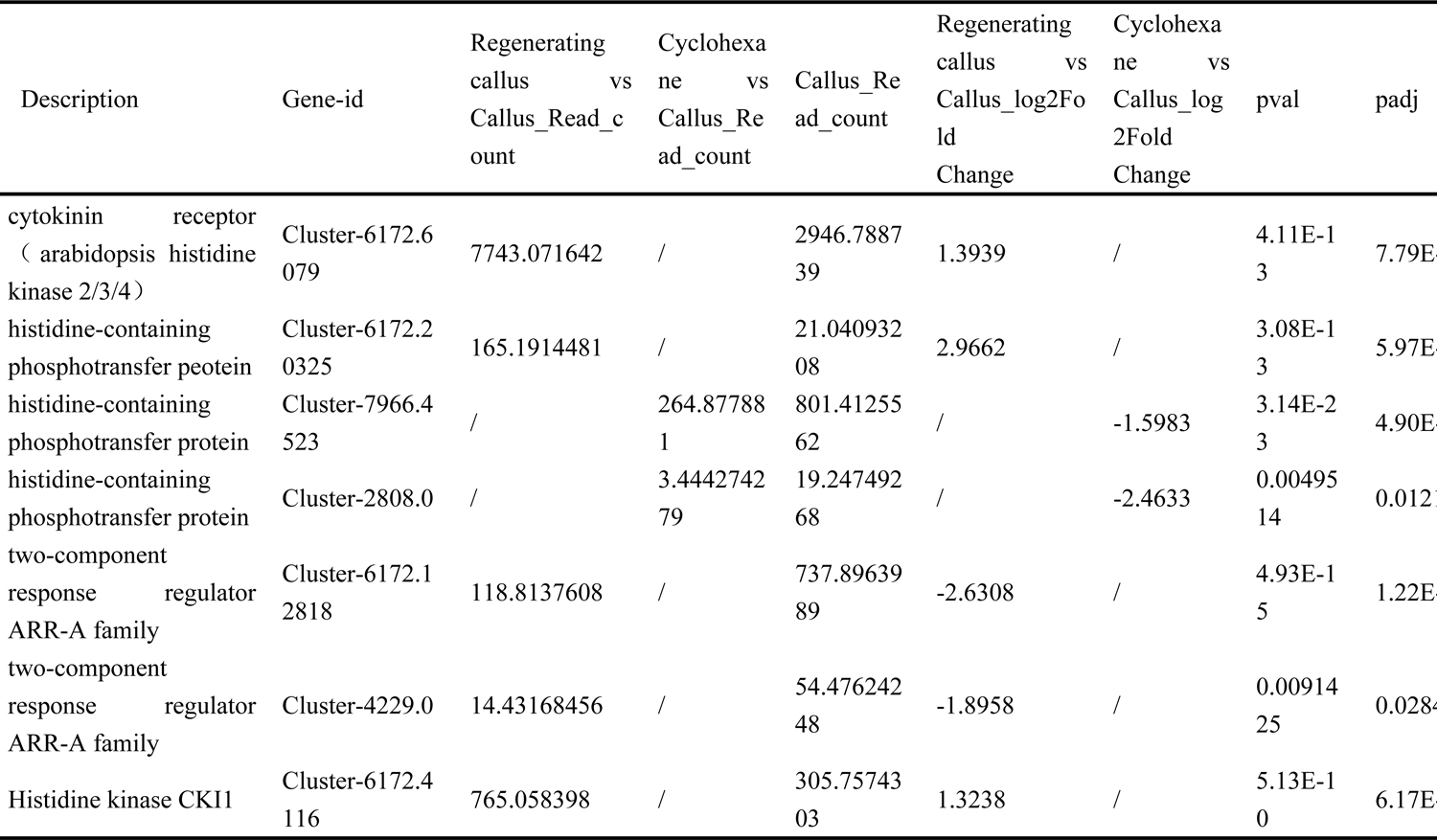
Gene expression in plant regeneration of Cytokine

Secondly, we studied the expression of genes related to CTK signal (Fig. 8b). The gene regulation in regeneration and treated with cyclohexane is different. The CRE1 has been up-regulated in the regenerating callus, and that has been down-regulated in callus treated with cyclohexane.

Thirdly, the expression of genes related to brassionosteroid signal has been investigated. In the brassionosteroid signal pathway, the expression of brassinazole-resistant1/2 (BZR1/2) has been down-regulated in callus treated with cyclohexane and the regenerating callus (Table 4). In the brassionosteroid signal pathway, the expression of BZR1/2 has been down-regulated in in callus treated with cyclohexane and the regenerating callus.

**Table 4.** Gene expression in plant regeneration of Brassinosteroid

| Description | Gene-id | Regenerating callus vs Callus_Read_count | Cyclohexane vs Callus_Read_count | Callus_Read_count | Regenerating callus vs Callus_log2Fold Change | Cyclohexane vs Callus_log2Fold Change | pval | padj |
| --- | --- | --- | --- | --- | --- | --- | --- | --- |
| BRI1 kinase inhibitor 1 | Cluster-6172.8113 | 291.510962 | / | 769.9837 | -1.4001 | / | 3.62E-07 | 2.73E-07 |
| brassinosteroid resistant 1/2 | Cluster-6172.9208 | 243.3127962 | / | 545.7915 | -1.1678 | / | 3.52E-07 | 2.66E-07 |
| brassinosteroid resistant 1/2 | Cluster-6401.0 | / | 43.03949153 | 146.1382 | / | -1.7659 | 9.00E-11 | 5.52E-11 |
| brassinosteroid resistant<br>1/2 | Cluster-6172.2<br>0298 | 33.13147023 | / | 156.7584 | -2.2381 | / | 5.50E-0<br>7 | 4.01E |
| cyclin D3 | Cluster-6172.6<br>746 | 932.3401808 | / | 2811.633 | -1.5916 | / | 3.80E-2<br>1 | 1.91E |

Moreover, the expression of genes related to ethylene signal has been investigated (Fig. 8c). The expression of ETR and EBF1/2 has been up-regualted in callus treated with cyclohexane and the regenerating callus. transcription factor MYC2(MYC2), plays a role in jasmonic acid signal pathway, has been up-regulated in both cyclohexane treatment and regenerating callus (Fig. 8d). There is no significant difference in gibberellin signal pathway during cyclohexane treatment (Fig. 8e).

## Discussion

In line with previous studies, we established an effective way to *in vitro* callus regeneration in duckweed. Interestingly, we found that one regenerating callus promoted another callus to regenerate. Genomes and transcriptome sequencing (especially plant hormones) and volatile substances were studied to reveal the molecule framework of plant regeneration in duckweed.

Plant hormones played a crucial role during callus regeneration^1^. Through our study, we hope to have a deeper understanding of the regulatory mechanism of callus regeneration. Callus were induced by auxin, similar as lateral root primordium^17–19^. In Arabidopsis, the callus tissue formed root stem cell niche, by regulation the expression of root stem cell regulators, including WOX^20–23^. According to our results, ARF, AUX/IAA, GH3, ARF1, SAUG and other response factor have been down-regulated significantly during the callus redifferentiation (Fig. 3). In the auxin signaling pathway, the interaction between ARF and AUX/IAA could regulate the genes expression of auxin early response. Moreover, ERF3, WOX11 and ZFP were found to be related to the ARs, LRs and RHs of initiation in *Spirodela*^8^, which might lead associated to the regeneration in duckweed.

Cytokinins and auxin have synergistic or antagonistic interactions with each other^24^. As a phytohormone, cytokinin could not only control key aspects of environmental responses, such as biotic and abiotic stress resopnses, but also regulate various developmental processes including cell proliferation, leaf formation, and root formation and growth^25–26^. Cytokinins promoted plant regeneration by regulating the generation of somatic embryogenesis in *Fumariaceae* and Rice^27–28^. In this study, cytokinin receptor CRE1, CKI1 and transfer protein of histidine phosphate AHP were up-regulated, during the expression of negative feedback regulator A-ARR was down-regulated in callus regeneration (Fig. 4). And the expreission of cytokinins synthesis was up-regulated, thereby promoting the differentiation of shoots. The transcriptome analysis suggested the similar result with Arabidopsis, giving evidence that the regulation of auxin and cytokinins leads to regeneration. Besides, plant regeneration has been regulated by other hormones^29^. In our results, we found that gibberellin, jasmonic acid and increased significantly, while genes related to gibberellin and brassinolide were down-regulated during callus regenerating (Fig. 8).

Plants release VOCs to the environment to affect their own or other biological life processes in the process of plants growth and development. This phenomenon was called allelopathy^30^. Plants in different growing environments, such as biological stress or abiotic stress, might release different VOCs to improve their resistance to external interference^31–33^. In previous studies, VOCs have been shown to mediate cell to cell communication, there by leading to stress responses in plants^34^. In our study, 11 kinds of specific VOCs have been increased during callus regenerating. Among them, cyclohexane could significantly promote the regeneration of callus in 16 days (Fig. 6).

Here, the regulation of gene expression related to hormone in callus treated with cyclohexane, which promoted regeneration, suggested the role of auxin during regeneration. AUX/IAA and GH3 has been down regulated in both callus treated with cyclohexane, which is similar with that in the regenerating callus (Fig. 8). And adventitious root initiation and enlongation has been promoted by AUX/IAA^8^. Interestingly, the root formation has been enhanced significantly by cyclohexane treatment (Fig. 6).

Altogether, we propose a hypothesis how callus regenerate in duckweed. Based on the DEGs in regenerating callus, we proposed molecular regulation on plant hormone. Also, our study provides candidates for evaluating the involvement of VOCs during duckweed regeneration, especially the enhancement of regeneration by cyclohexane. It also provides a resource for comparative transcriptome analysis of plant regeneration in other species.

It was indiciatied that VOCs might played an crucial role in the process of plant regeneration. It also makes clear that allelopathy does affect plant growth and development.

## Materials and methods

### Plant material and *in vitro* establishment and cyclohexane treatment

Lemna turionifera used in the experiment were collected from a lake in Tianjin, China. Duckweed was cultured in the liquid medium descripted as Wang et al. and Yang et al^35–36^. The duckweed was cultured aseptically in the liquid medium. Fully expanded fronds were selected as explant for callus induction. The rhizoid was removed, and the frond was scratched for callus induction. The induction medium was B5 solid medium, which was designed by Gamborg for soybeans tissue culturein 1968^37^. The induction medium contained plant hormones 15 mg/l dicamba, 3.5 mg/l 2, 4-D, 6-BA 2mg/l and 1.5% sucrose. The pH of medium was adjusted to 6.2-6.4 and then it was sterilizated at 121℃ for 20 minutes. The tissue was cultured in an incubator with a light cycle of 23 ± 2 ℃, 16 hours of light and 8 hours of darkness. After 4-5 weeks of induction, the duckweed explants developed into callus through dedifferentiation. After 2-3 weeks of induction, calli formated. The calli were transferred to the subculture medium. Subculture medium contains B5 medium, 10 mg/L 4-chlorophenoxyacetic acid (CPA) and 2 mg/L 2ip. In order to keep the callus with better morphology and activity, a new subculture medium was replaced every two weeks. Callus was transferred to the regeneration medium for duckweed regeneration. The regeneration medium contains B5 medium, 1 mM serine, and 1.5% sucrose. After 2 or 3 weeks, the callus redifferentiated and regenerated.

When 3 days culture in B5 subculture medium, the calli were cultured in B5 medium with 20 ml cyclohexane in a large airtight beaker. Each day open the sealing device regularly to change the air in the beaker. And replace with a new cyclohexane every two days.

### The Co-culture of regenerating callus and callus

The callus was cultured on subculture medium for more than two weeks for subsequent experiments. Callus and regenerating callus in the same growth condition were placed in B5 medium (containing 1.5% sucrose) respectively. For fumigate, the regenerating callus and callus were placed together in a closed environment for co-culture described as Fig. 9a.

**Fig. 9.**
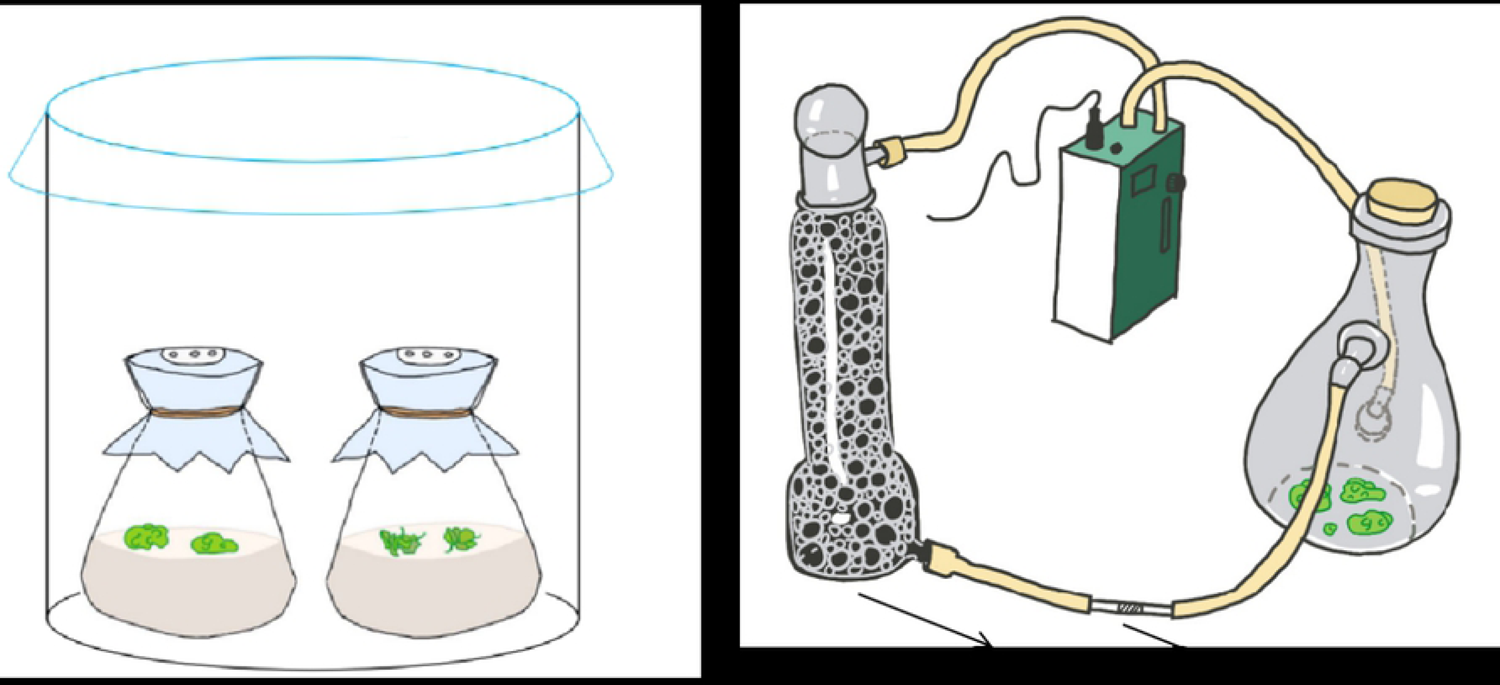
System of co-culture and dynamic headspace air-circulation. a The callus and the regenerating callus of duckweed were fumigating treatment. **b** Collection of VOCs from plant tissue. (i) Activated carbon. (ii) Adsorption tube.

### VOCs Collection and analysis

Shown as Fig. 9b, the VOCs released from callus and regenerating duckweed were collected using the dynamic headspace air-circulation method described by Zuo et al. (2018)^38^. There were 3 conical flasks of callus or regenerating callus for each group. The chemical composition analysis of VOCs was performed by thermal-desorption system/gas chromatography/mass spectrum (TDS/GC/MS). And the GC/MS data was studied in NIST/EPA/NIH Mass Spectral Library (NIST 08) (National Institute of Standards and Technology, MD, USA).

### RNA isolation, quantification, and sequencing

RNA degradation and contamination on 1% agarose gel were detected, and the quality of the samples was qualified. RNA purity was checked using the NanoPhotometer® spectrophotometer (IMPLEN, CA, USA). RNA concentration was measured using Qubit® RNA Assay Kit in Qubit® 2.0 Flurometer (Life Technologies, CA, USA). And then, RNA integrity was assessed using the RNA Nano 6000 Assay Kit of the Agilent Bioanalyzer 2100 system (Agilent Technologies, CA, USA).

### Sequencing data filtering and transcript assembly

Image data from sequencing fragments measured by high-throughput sequencers are transformed into sequence data (reads) by CASAVA base recognition. The raw data obtained from sequencing included a small number of reads with sequencing adaptors or low sequencing quality. The filtering contents were followed as our previous study: Removed adapters; Removed reads whose proportion of N is greater than 10%; Remove low-quality reads^6^. The clean reads were assembled by the trinity de novo assembly program with min_kmer_cov set to 2 by default, otherwise it was set to default^39^. Overall, a reference sequence, with an average length of 1928 bp and a total length of 282527137 bp, was obtained for subsequent analysis.

## Data analysis

The experiment were repeated for at least triplicate independent experiments. Analysis of variance (ANOVA) method and SPSS software (IBM SPSS Statistics, Version 20) were applied to compare the statistical significances. Significant difference in experiment was indicated by asterisks (*P < 0.05, **P < 0.01). And standard deviations were shown by error bar. The graphs in this studies were made using Origin 9.0 (Origin Lab, USA).

## Data availability

All data included in this study are available upon request by contact with the corresponding author.

## Author contributions

Sun, J.S. provided resource support and orchestrated the arrangement. Yang, L. designed the experiment,, analyzed the data and wrote the manuscrip. Sun, J.G. finished the experiment and counted the experimental results. Yao, J. supervised the experimental project and the subsequent editing of the manuscript. Wang, Y. Y. completed the data statistics and forms. Yan, C.Y. completed the chart drawing and part of the experiment. Wu, J.Y. adjusted the overall format of the manuscript and completed basic experiments. Ren, Q.T. cultured the experimental plants. Zhao, L. provided technical assistance.

